# Haplotype mapping uncovers unexplored variation in wild and domesticated soybean at the major protein locus cqProt-003

**DOI:** 10.1101/2021.10.12.464159

**Authors:** Jacob I. Marsh, Haifei Hu, Jakob Petereit, Philipp E. Bayer, Babu Valliyodan, Jacqueline Batley, Henry T. Nguyen, David Edwards

## Abstract

Here, we present association and linkage analysis of 985 wild, landrace and cultivar soybean accessions in a pan genomic dataset to characterize the major high-protein/low-oil associated locus cqProt-003 located on chromosome 20. A significant trait associated region within a 173 kb linkage block was identified and variants in the region were characterised, identifying 34 high confidence SNPs, 4 insertions, 1 deletion and a larger 304 bp structural variant in the high-protein haplotype. Trinucleotide tandem repeats of variable length present in the third exon of gene *20G085100* are strongly correlated with the high-protein phenotype and likely represent causal variation. Structural variation has previously been found in the same gene, for which we report the global distribution of the 304bp deletion and have identified additional nested variation present in high-protein individuals. Mapping variation at the cqProt-003 locus across demographic groups suggests that the high-protein haplotype is common in wild accessions (94.7%), rare in landraces (10.6%) and near absent in cultivated breeding pools (4.1%), suggesting its decrease in frequency primarily correlates with domestication and continued during subsequent improvement. However, the variation that has persisted in under-utilized wild and landrace populations holds high breeding potential for breeders willing to forego seed oil to maximise protein content. The results of this study include the identification of distinct haplotype structures within the high-protein population, and a broad characterization of the genomic context and linkage patterns of cqProt-003 across global populations, supporting future functional characterisation and modification.

**Key message:** The major soy protein QTL, cqProt-003, was analysed for haplotype diversity and global distribution, results indicate 304bp deletion and variable tandem repeats in protein coding regions are likely causal candidates.

## Introduction

Shifting climatic and ecological conditions threaten global food security at a time when the growing human population requires crop yields to increase an estimated +50% to +110% by 2050 (Alexandratos and Bruinsma 2012; Ray et al. 2013; Tilman et al. 2011; van Dijk et al. 2021). Domestication and improvement of major crops has led to genetic bottlenecks and reduced diversity due to strong selection for agronomic traits, especially in self-pollinating plant species such as soybean (*Glycine max* (L.) Merr.) (Hyten et al. 2006). Whilst intensive breeding efforts have increased crop productivity, it has left the regions of the soybean genome under selection with low genetic diversity in some modern breeding populations (Zhao et al. 2015). The lack of variation in these regions is particularly concerning, as they have proven to play important roles in plant function or morphology, and yet there is limited allelic variation remaining in modern lines for trait expansion and adaptation. When dissecting the genomic regions underlying agronomic traits it is important to look beyond traditional breeding populations and capture the full range of potential diversity. Fortunately in soybean, ancestral diversity persists in the wild progenitor *Glycine soja* (Siebold & Zucc.) and exotic landrace populations that still harbour genomic variation at loci that are of value to breeders (Kofsky et al. 2018; Zhang et al. 2017).

Broad resequencing of soybean germplasm has provided a wealth of data for characterising genetic diversity and for identifying genomic variation underlying agronomic traits (Fang et al. 2017; Torkamaneh et al. 2021a; Valliyodan et al. 2021; Zhou et al. 2015). In addition, the development of pangenomic datasets that increase variant and marker data accessibility across germplasm collections offers novel opportunities for plant breeding studies by providing the entire genetic content of soybean, including small and large structural variations, across global populations (Bayer et al. 2021; Liu et al. 2020; Torkamaneh et al. 2021b). Several studies have mapped important major effect genomic loci; for example disease resistance (Chang et al. 2016), SCN resistance (Patil et al. 2019); mycorrhizal colonization (Pawlowski et al. 2020); salt tolerance (Do et al. 2019); seed composition (Patil et al. 2018); and descriptive traits such as flower and pubescence colour (Bandillo et al. 2017). Genetic approaches incorporating whole-genome information with breeder friendly phenotyping (Reynolds et al. 2020) have allowed precise dissection of the genetic architecture of complex traits including seed composition in soybean. However, comprehensive bioinformatic analysis of key traits, such as seed compositional traits, are needed to capitalize on this fast accumulating wealth of soybean data for agronomic gain (Marsh et al. 2021).

Protein content in soybean is associated with a major quantitative trait locus (QTL); cqProt-003 on chromosome 20, that underwent a genetic bottleneck during domestication with selection for oil, at the cost of fixing a low-protein phenotype across modern breeding populations (Brzostowski et al. 2017). The region associated with cqProt-003 was first detected as a result of QTL mapping in an experimental population derived from crossing a domesticated *G. max* line with a wild *G. soja* individual (Diers et al. 1992). This seed protein QTL was later genetically mapped to a 3 cM region on Linkage Group I (Nichols et al. 2006), before being located to an 8.4 Mb genomic region on chromosome 20 (Bolon et al. 2010). Attempts over the following decade to refine the genomic locus responsible for seed composition variability have produced conflicting reports, in part due to the extensive linkage present across the region (Bandillo et al. 2015; Cao et al. 2017; Hwang et al. 2014; Vaughn et al. 2014). It has been suggested that a 321 bp structural variant in an exonic region of the gene *20G085100* (CCT motif family protein) may be responsible for the phenotypic effect (Fliege 2019). However, questions remain as to whether other variation is present in the high protein haplotype, and the genomic context and the distribution of the high protein haplotype across diverse international lines are largely uncharacterized.

Here we take a comprehensive approach to characterizing genomic variation at the cqProt003 locus using recently published high quality pangenomic datasets from wild and domesticated soybean populations. This study aims to characterize the haplotype variation present within this region and evaluate the potential effects of allelic variants that may contribute to the high-protein phenotype. This will provide breeders and researchers with a detailed map of untapped variation in the cqProt-003 region that can serve as a roadmap for crop improvement.

## Materials and methods

### Genotyping and additional genomic resources

The SNP matrix was generated from 985 of the soybean lines in the previous soybean pangenome study (Bayer et al. 2021). Briefly, adapters of raw reads were removed using Trimmomatic v0.36 (Bolger et al. 2014). Clean reads were mapped to the Wm82.a4.v1 (Valliyodan et al. 2019) reference genome using BWA-MEM v0.7.17 with default settings (Li 2013) and duplicates removed by Picard tools (http://broadinstitute.github.io/picard/). Reads were realigned by GATK (McKenna et al. 2010) v3.8-1-0 RealignerTargetCreator and IndelRealigner, followed by variant calling using GATK HaplotypeCaller. Then, indels and SNPs residing on unassembled scaffolds were split into separate files. The resulting SNPs were filtered (QD< 2.0 || MQ < 40.0 || FS > 60.0 || QUAL <60.0 || MQrankSum < −12.5 || ReadPosRankSum < −8.0) to remove low-quality SNPs. SNP loci with more than 10% missing data and less than 1% minor allele frequency were filtered out using VCFtools v3.0 (Danecek et al. 2011), resulting in 544,997 high quality SNPs for chromosome 20. The filtered dataset included Phasing and imputation of missing genotypes was conducted using BEAGLE v5.1 (Browning and Browning 2007). Variant annotation was predicted using SnpEff v3.0 (Cingolani et al. 2012). Nucleotide diversity (π) and divergence (dXY) was calculated using pixy v1.0.4.beta1 using all invariant sites (Korunes and Samuk 2021), with all sites filtered to max-missing 0.9 and variant sites only filtered with --minQ 30 using VCFtools v3.0 (Danecek et al. 2011).

### Insertions, deletions and structural variation

Insertions and deletions were recoded to biallelic sites to enable straightforward linkage calculation with SNPs. Insertions were split from deletions based on the relative lengths of the alternate alleles compared to the reference using BCFtools norm -m (Li et al. 2009). The insertion and deletion files were then filtered (--minQ 30 --max-missing 0.9 --non-ref-af-any 0.01) with VCFtools v3.0 (Danecek et al. 2011). Multi-allelic records were then re-merged for insertions and deletions before being recoded to biallelic sites with all variant alleles of the same type (i.e. insertion or deletion) represented by a single proxy alternate polymorphism (e.g. Adenine), while the reference allele was also recoded to a different proxy allele (e.g. Tyrosine). For chromosome 20, 35,826 Insertions and 55,512 deletions were kept following filtering and recoding. Recoded biallelic insertions and deletions were then merged with filtered SNPs in separate files in order to preserve 4540 multiallelic sites with both insertions and deletions. Presence and absence for the 304bp structural variant was classified from significant drops (8-50X to 0-1X) in per-base coverage in the BAM file across the *20G085100* genic region using mosdepth v0.2.6 (-b 1 -Q 10) (Pedersen and Quinlan 2018).

### Phenotype collection and Metadata

The protein and oil phenotypic data used in this study were obtained from the USDA-Soybean Germplasm Collection general evaluation trials which includes morphological, agronomic and seed composition data sets. These field evaluations were conducted at different locations based on the maturity group where some of them were grown for several years and the total protein and oil concentration were measured using various methods. The protein and oil concentration measurements of the soybean accessions with yellow seed coat were conducted using the near-infrared reflectance method on whole seed sample. The dark or pigmented seed coated soybean samples were analysed for total protein content using the Kjeldahl method and the seed oil content using the Butt method (Bandillo et al. 2015). There are several reports using these data sets for genotype-phenotype association studies and identification of oil and protein content genes or QTL in soybean (Bandillo et al. 2015; Hwang et al. 2014; Jarquin et al. 2016; Vaughn et al. 2014). List of accessions used in this study with the available protein and oil data is available in Table S6.

### Genome-Wide Association Analysis

Association analysis was performed using the FarmCPU (Liu et al. 2016) and MLM (Price et al. 2006) methods implemented in rMVP (Yin et al. 2021). The GWAS included data from all 985 lines for protein, and the subset with data available of 945 for oil (Table S6) with imputed and unimputed data (Table S1, Table S3). Population structure was controlled using the first three PCAs based on the marker data. The significance threshold was determined by 0.05/marker size.

### Linkage disequilibrium analysis

The linkage disequilibrium heat map was generated using LDBlockShow (Dong et al. 2020) for R^2^ and D’ with -block type set to the Gabriel method for the D’ figure (Gabriel et al. 2002), the input for the Manhattan plot was the protein GWAS FarmCPU results from the filtered, unimputed SNP data. The approach used for the detailed haplotype block information was the confidence interval definition (Gabriel et al. 2002) implemented in PLINK1.9 (Purcell et al. 2007) with default parameters. Pairwise R^2^, D’ and allele frequency values were obtained for sites using PLINK1.9 (Purcell et al. 2007) by first using --show-tags with --tag-r2 0.9 to identify sites in linkage with the GWAS-SNP, before generating detailed information with this list of sites with --r2 dprime. Site missingness information was generated using VCFtools v3.0 (Danecek et al. 2011).

To avoid false positive correlations, linkage analysis was conducted on filtered, unimputed SNPs with recoded biallelic insertions and deletions. Linkage requirements to include insertions/deletions in the cqProt-003 haplotype were relaxed to R^2^ > 0.85 to compensate for the loss of information from recoding with merged variants. For the 304bp structural variant, individuals with presence/absence were recoded the same way for linkage testing, and the relatively higher impact larger variations can have compared to point mutations. Linkage between SNPs and insertions/deletions was testing between the presence of an insertion/deletion of any length compared to the reference allele, rather than specific variants for multiallelic sites.

### Site-centric haplotyping

Site-centric haplotyping was conducted using HaplotypeMiner software (Tardivel et al. 2019). The genomic input was generated using the filtered, imputed SNP dataset converted to hapmap format using Tassel 5 (Bradbury et al. 2007), with heterozygotes set to missing, from which only sites within the 173kb (31,604,127 to 31,777,346) region were retained. The R2 measure was ‘r2vs’ with a kinship file generated using the Endelman & Jannink (Endelman and Jannink 2012) method implemented in Tassel 5 (Bradbury et al. 2007) and a structure file generated using fastStructure (Raj et al. 2014) with optimal K = 4, --prior = simple. The cluster R2 was set to ‘r2’ only to reduce the computational load. The ‘gene’ region provided was the start and end position of the previously determined haplotype block defined that contains the GWAS-SNP; start: 31621924 end: 31644477. The following parameters were applied: max_missing_threshold = 0.6, max_het_threshold = 0.3, min_alt_threshold = 0.01, min_allele_count = 4, marker_independence_threshold = 0.9, with a range of cluster thresholds tested using the clustree R package (Zappia and Oshlack 2018). Segmentation between cluster groups was assessed by performing PCA implemented in the factoextra R package which provided input for UMAP (McInnes et al. 2018) with the following parameters: -pc 7 -nn 15 -md 0.001.

Linkage of other sites with the representative marker SNPs that define the haplotypes was conducted independently using PLINK1.9 (Purcell et al. 2007) with --tag-r2 0.9 on the unimputed dataset. Welch’s independent two-sample t-test assuming unequal variance (Welch 1947) was used to test for significant differences of protein content between individuals across haplotype groups and marker groups. Six marker groups that were co-inherited in every haplotype population with another representative marker were removed to reduce redundancy, keeping only one representative marker; a further six marker groups were removed which either included only five or fewer SNPs or differed from another marker group by only one SNP. Graphical summaries of the haplotype populations and marker groups with phenotype information was conducted using ggplot2 in R v4.04 (R Core Team 2021).

## Results

### Phenotype association analysis

A genome-wide association study (GWAS) for protein content was conducted on chromosome 20 using 985 accessions, including 131 wild lines, 708 landraces, 44 old cultivars and 102 modern cultivars, for which phenotypic data was available from the UWA SoyPan dataset (soybase.org). GWAS was tested using different combinations of unfiltered, filtered, unimputed and imputed data (Figure S1, Figure S2, Figure S3, Table S1). The results using the FarmCPU method and the unimputed dataset indicate a highly significant site associated with protein content located at position 31,649,589 in the Wm82.a4.v1 reference genome (Valliyodan et al. 2019)) (Table S1); however, a site 17 kb upstream (31,632,556) was detected when using imputed data (Figure S4, Table S1). Given the 9.3% missing variant information from lines that did not align at position 31,649,589 (Figure S5, Table S2), the SNP at 31,632,556 (hereafter referred to as “GWAS-SNP”) was taken as the most confident GWAS result using this method. Using the MLM method, no sites registered above the significance threshold (p<1.0E^−08^); the most significant site was at 31,604,127 bp with a p-value of 3.15E-07, with 13 of the top 20 sites all within the 104 kb range of 31,604,126-31,708,981 bp (Table S1). GWAS for oil content was conducted on the same dataset using the FarmCPU method and 945 individuals (Table S3). A significant site corresponding to decreased seed oil (p = 3.4e-09) was detected at 31,687,912 bp, at which the 183 individuals with the alternate allele displayed a mean oil content 32.7% lower (6.1g oil per 100g seed weight) than the 743 individuals conforming to the reference.

### Delimiting the cqProt-003 region

The chromosomal region associated with the protein phenotype was defined from linkage disequilibrium surrounding the GWAS-SNP. Significant, though non-continuous linkage, defined using R^2^, is apparent in large blocks across a wide 300 kb region (Fig. 1), with a block of LD surrounding the GWAS-SNP from approximately 31,600,000-31,800,000. Linkage blocks in the region defined using the confidence interval approach are relatively continuous from 31,601,763-31,732,110 bp, becoming more disconnected in adjacent regions (Figure S6, Table S4).

**Fig. 1.**
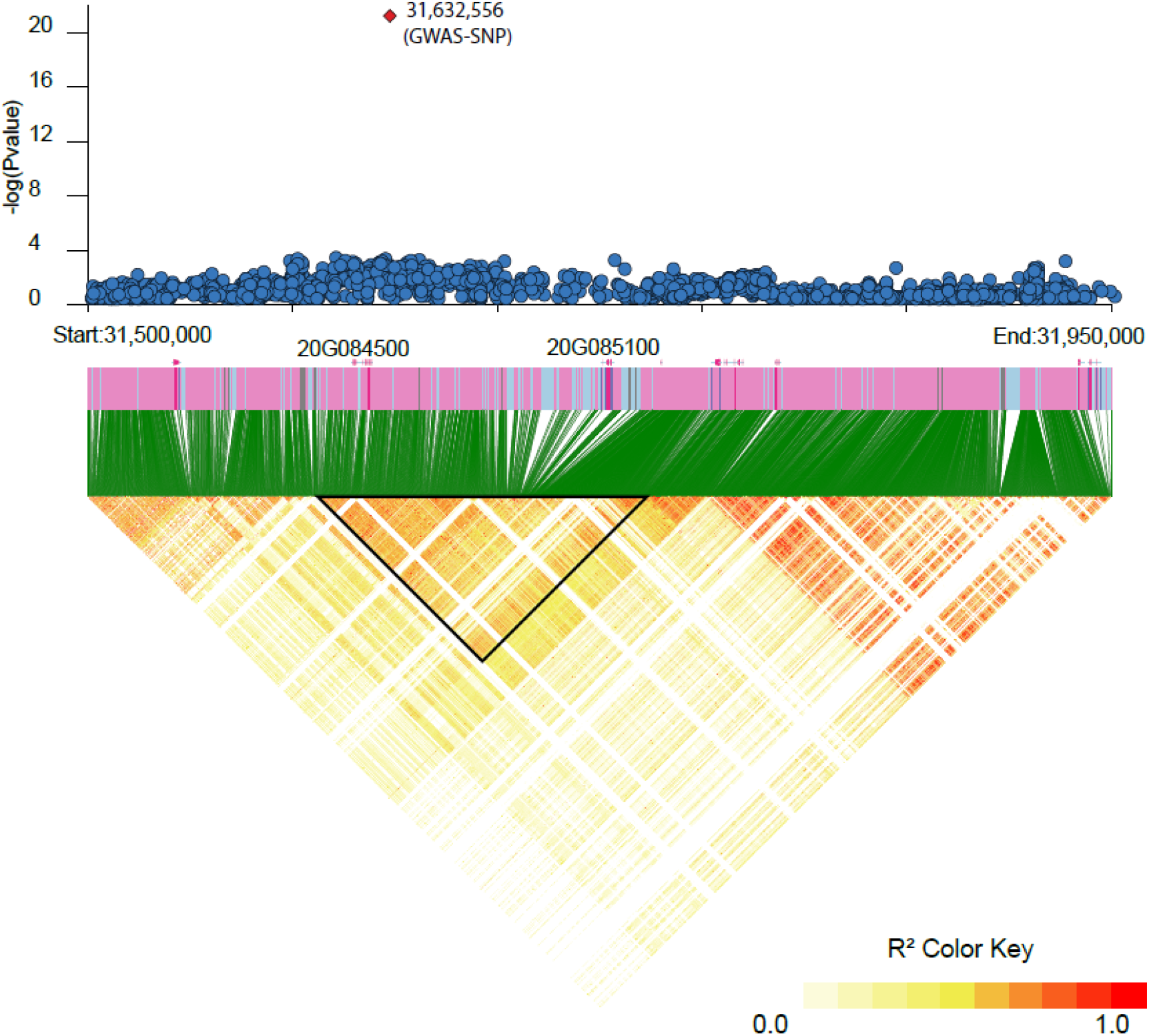
**Linkage disequilibrium (R^2^) between SNPs** across 450Kb region surrounding the significant protein GWAS hit at 31,632,556. The black triangle represents the 173kb cqProt-003 region defined from decay of linkage with the GWAS-SNP only.

Linkage with the GWAS-SNP assessed using R^2^, remained relatively high at long distances, with a value above 0.7 beyond 1 Mbp downstream (Figure S7), though R^2^ linkage is only present above 0.8 across a 330 kb region from 31,590,913 to 31,920,542 bp, and above 0.91 across a 173 kb region from 31,604,127 to 31,777,346 bp. D’ for sites that have an R^2^ greater than 0.7 decay precipitously as distance increases from the site, before D’ plateaus below 0.967 beyond the same range designated by the 0.91 R^2^ cut-off: 31,604,127 to 31,777,346 bp (Figure S8). The 173 kb linkage region spans seven significant blocks (>10 SNPs) of linkage defined by the confidence interval approach (Gabriel et al. 2002) (Table S4). Pairwise linkage between SNPs was extensive, with a distinct stratum of very strongly linked SNPs (R^2^ ≥ 0.91) across the region (Figure S9). It is notable that rates of site missingness spiked flanking the region, especially upstream in the range from position 31,593,149 to 31,601,877 bp, and downstream of the *20G085100* gene at 31732565 to 31740664 bp within which all sites had missing reads with a mean above 5% (Figure S5), potentially indicating structural diversity, problematic read alignments or mis-assembly in the Wm82.a4 reference.

### Patterns of linkage present within the region

Gene-centric haplotyping was conducted with SNPs in the 173 kb region, producing distinct groups of internally linked markers as well as groups of individuals with different haplotype combinations of these marker groups. The haplotyping resulted in nine groups of individuals (A-G combinations) that contained different combinations of six groups of markers (Fig. 2), which include 370 SNPs that are in tight linkage with one of the different representative sites (M01-M06) (Fig. 3, Figure S10, Figure S11, Figure S12, Table S5, Table S6). Comparing the phenotype score between individuals from different haplotype groups indicates a significant elevation of protein content (p = 7.47e^−14^), with a mean increase for landrace individuals of 3.32 g/100 g seed weight individuals with the alternate allele for marker group M01; conversely, oil content displays a mean decrease of 2.66 g/100 g seed weight (Tables S7, Table S8). These high protein/low oil haplotypes (B-G) are comprised mostly of wild and landrace individuals, compared to haplotype group A which is predominantly domesticated (Fig. 4). Wild individuals display consistently lower oil content with highly variable protein content; however landrace individuals display a high degree of oil variability within and between haplotypes (Fig. 4). Notably, haplotype group C represents an exclusively landrace population (n= 21, μProtein = 48.762, S^2^ = 6.902) that remains consistent with the high protein phenotype seen in other haplotype populations that contain a mixture of wild and landrace individuals (Fig. 4, Table S7).

**Fig. 2.**
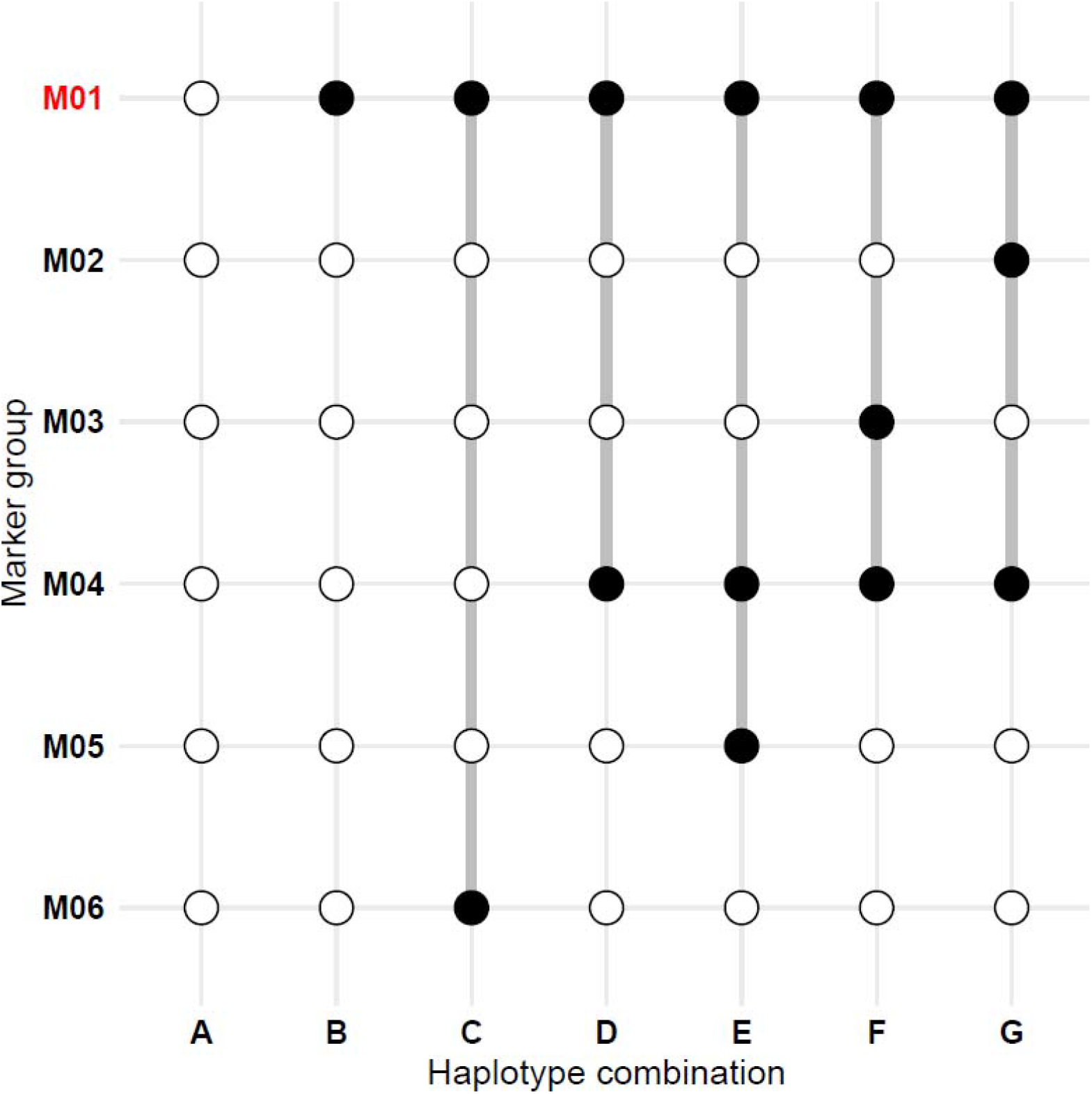
**Region-centric haplotypes for 173kb region** indicating alternate alleles (black dots) for clustered markers (M01-M06) that define unique haplotype combinations (A-G) in the 173kb region of interest; white dots indicate reference alleles present. The protein-associated haplotype, M01, is highlighted in red.

**Fig. 3.**
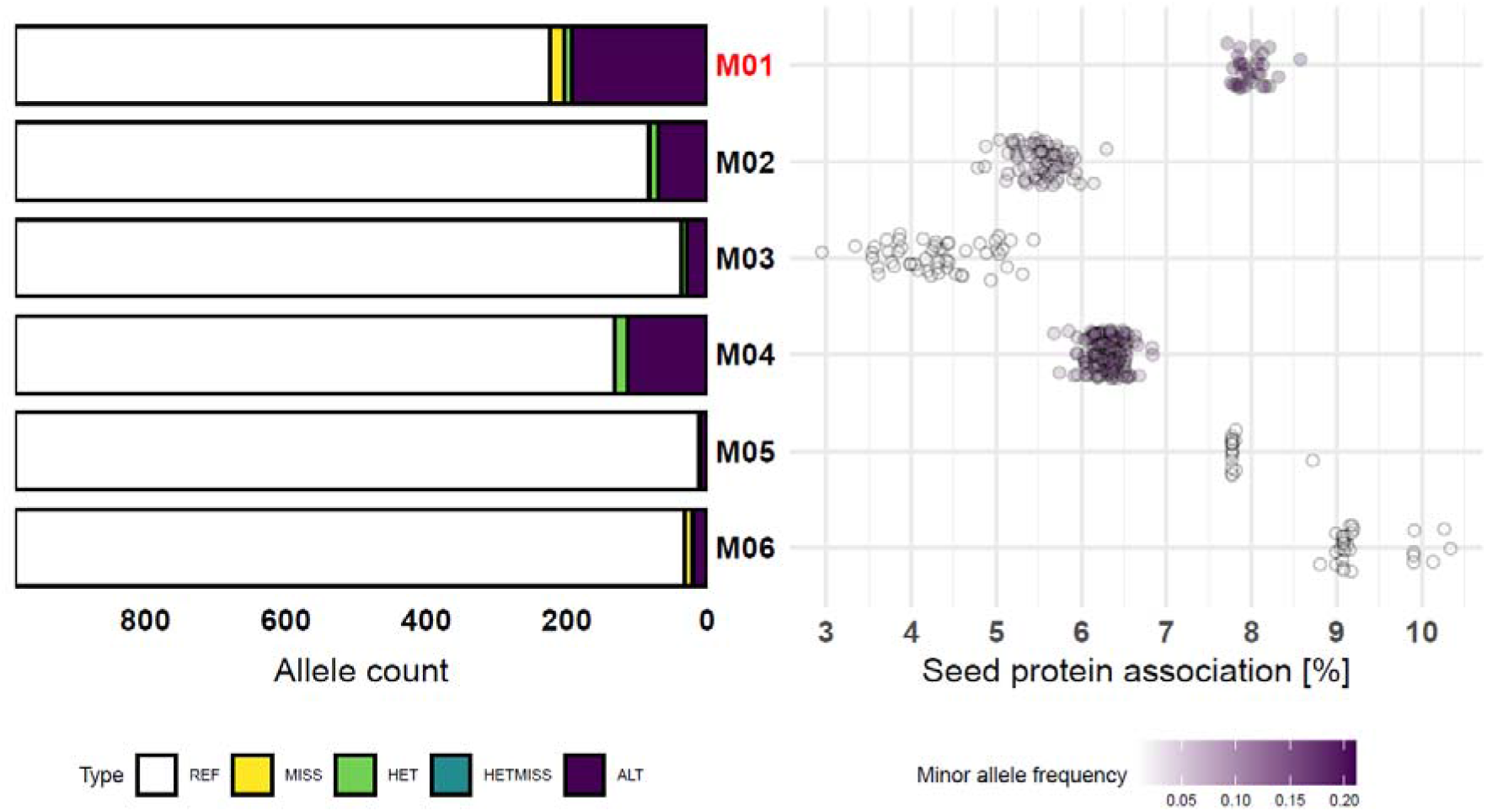
Allele frequencies for the representative markers (left) and summary of supporting markers in each group (right). Marked in red is the marker group containing candidate variants for the high protein haplotype (M01). Each dot represents the average protein difference between individuals who have the alternate vs reference allele for a single SNP within a given marker group, coloured white-purple by that site’s alternate allele frequency. All alternate alleles depicted are positively associated with seed protein, compared to the reference allele. REF refers to the frequency of homozygous reference alleles at the marker groups’ representative site, similarly for homozygous missing (MISS), heterozygous (HET), sites with one missing allele (HETMISS) and homozygous alternate alleles (ALT).

**Fig. 4.**
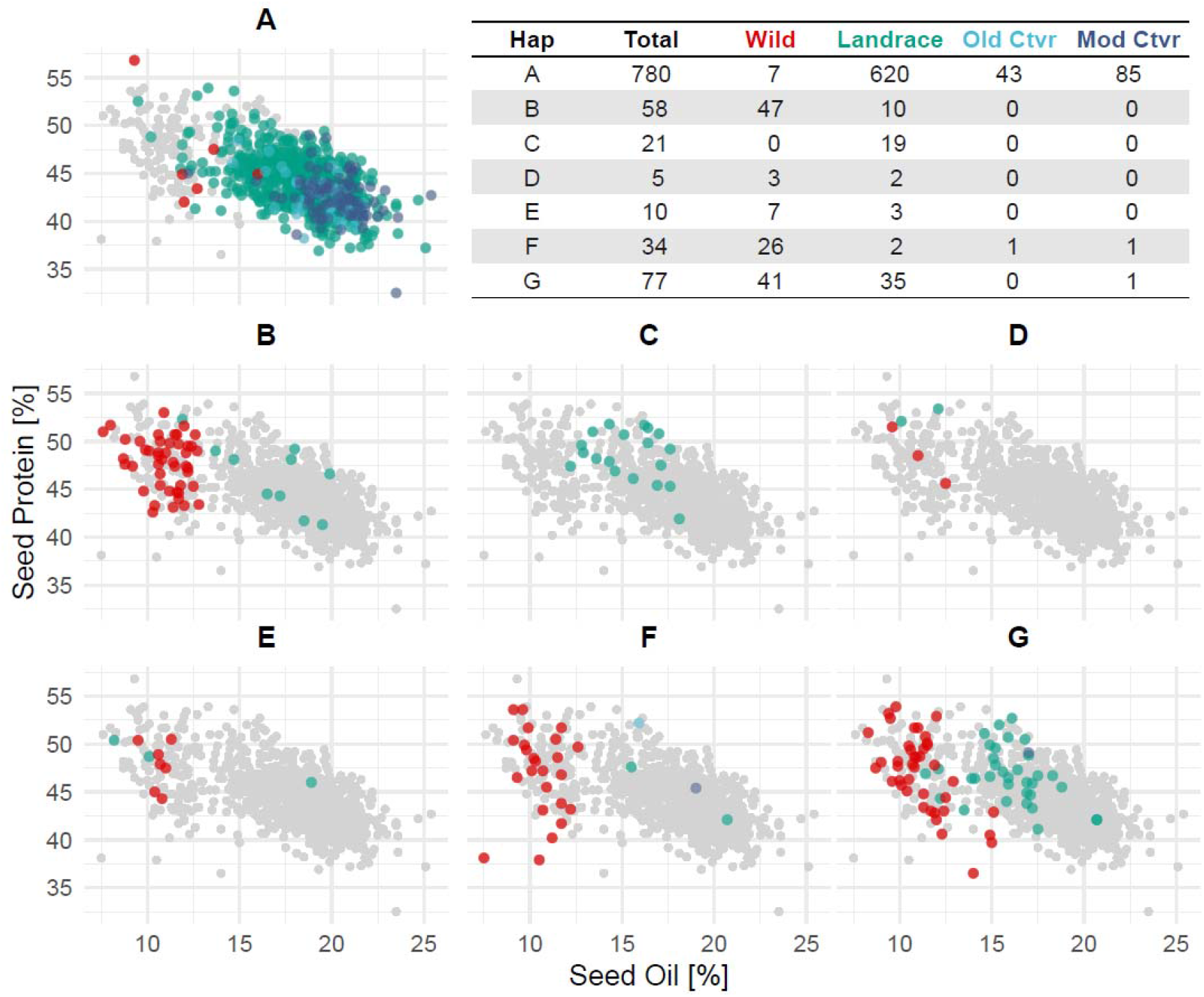
Phenotype associations of haplotype combinations (A-I) with population breakdown. Each dot represents an individual possessing a given haplotype population (A-G), coloured by level of domestication: wild, landrace, old cultivar (‘Old Ctvr’) and modern cultivar (‘Mod Ctvr’); grey points include all individuals for reference.

Genomic variation across the region is primarily present in individuals with the alternate allele in the M01 marker group (Fig. 2, Fig. 3). In the 173 kb genomic region, nucleotide diversity (π) is 3.85E^−3^ among individuals with the alternate allele for the M01 marker group, compared to π = 3.24E^−4^ with the reference allele. This represents a significant drop of diversity in the 173 kb region for the population with the M01 reference allele that possess an average diversity (π) of 1.74E^−3^ across the entire genome. In contrast, the population of individuals with the alternate allele at M01 has only slightly elevated genetic diversity in the region relative to the mean across the whole genome (π = 3.60E^3^). Increased genetic diversity in the region appears to be contained in the predominantly wild populations; of the alternate M01 haplotypes, only the landrace group C possesses lower levels of diversity consistent with the primarily domesticated haplotypes A and B (Table S7). Furthermore, divergence in the 173 kb region between groups A-C are far lower than groups D-G with the alternate M04 marker group (Figure S13, Figure S14), which includes 157 SNPs, though has a slightly lower association with seed protein than M01 (Fig. 3).

### Trinucleotide repeat expansion found in the high-protein haplotype

The haplotype of 40 variants in linkage with the GWAS-SNP (Table S9) was further characterised. Four insertions and a single deletion were also found to be in tight linkage (R^2^>0.85) with the GWAS-SNP, warranting their inclusion in the high protein haplotype (Table S9). Of the 39 small and medium sized variants classified as in the cqProt-003 haplotype, 33 were in intergenic regions and five were adjacent to genes (Table S9). There were no variants detected from the haplotype in coding regions of *20G084500* and *20G085250*. However, a multiallelic insertion at 31,727,019 bp is located in a protein-coding region, the first exon of *20G085100.1* (Table S10). The mutations at this locus appear to be predominantly trinucleotide tandem repeats of ‘CAA’, which are conservative in-frame insertions of Asparagine, however there are four individuals who reported either disruptive in-frame insertions or frameshift variants (Table 1). The phenotype scores for individuals that contain homozygous tri-allelic insertions are significantly elevated, with an average protein score increase of 7.9% compared to individuals without the insertion (Table 1), consistent with the cqProt-003 haplotype. The mean protein scores for individuals with each alternate sequence at the tandem repeat locus were all higher than the mean for individuals without an insertion at this site; however, sample sizes are insufficient for confident association testing between these alleles (Table 1). In landrace individuals, 82.3% with homozygous alternate insertions at tandem repeat locus possessed the seven or eight repeat allele, compared to 25.8% in wild individuals that are more diverse (Fig. 5).

**Table 1.** Summary of individuals with different alleles present at the trinucleotide repeat locus, 31,727,019, including sequence, annotation, individual count and mean protein content. The ‘other’ row refers to four individuals with insertions at this locus that did not fit the tandem repeat structure; ‘heterozygous repeats’ row refers to individuals that are heterozygous at the locus with two different insertions; ‘single repeat’ refers to individuals that are heterozygous with one copy of any insertion and one reference allele; ‘other’ refers to individuals that are homozygous for an insertion that is not a trinucleotide repeat of ‘CAA’; ‘missing’ refers to individuals with one or both missing alleles. Annotations include conservative in-frame insertion (CIF), disruptive in-frame insertion (DIF) and frame-shift variant (FSV).

| **31,727,019 sequence** | **Annotation** | **Individuals** | **Mean protein** |
| --- | --- | --- | --- |
| ref; C | N/A | 787 | 43.86 |
| CCAA | CIF | 2 | 49.85 |
| C('CAA'*2) | CIF | 10 | 46.67 |
| C('CAA'*3) | CIF | 8 | 45.64 |
| C('CAA'*4) | CIF | 21 | 46.75 |
| C('CAA'*5) | CIF | 13 | 48.19 |
| C('CAA'*6) | CIF | 24 | 47.19 |
| C('CAA'*7) | CIF | 46 | 46.68 |
| C('CAA'*8) | CIF | 30 | 48.97 |
| C('CAA'*9) | CIF | 1 | 50.20 |
| other | 2xDIF; 2x FSV | 4 | 44.78 |
| heterozygous repeats | N/A | 11 | 47.09 |
| single repeat | N/A | 15 | 47.03 |
| missing | N/A | 13 | 46.92 |
| **Total** | N/A | 985 | 44.54 |

**Figure 5.**
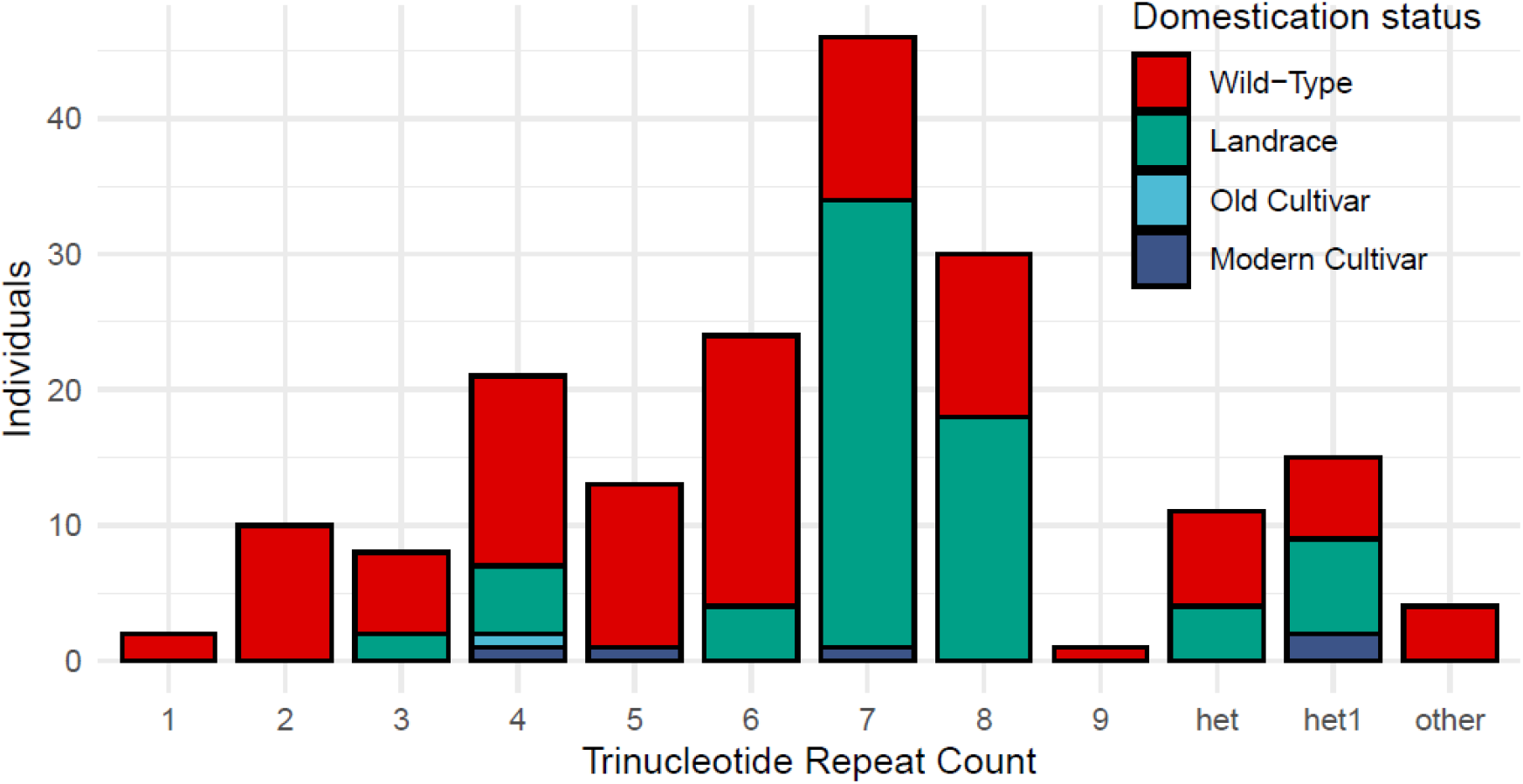
Frequency of different copy numbers of trinucleotide insertion alleles at the 31,727,019 locus, by domestication status. The ‘het’ column refers to individuals that are heterozygous at the locus with two different repeat insertions; ‘het1’ column refers to individuals that are heterozygous with a single copy of any repeat insertion and one reference allele; other refers to individuals that are homozygous for an insertion that is not a trinucleotide repeat of ‘CAA’.

### Trait associated structural variation

Further structural variation was found in the *20G085100* gene, with a 304 bp deletion (relative to Wm82). The deletion spans from 31,728,619 to 31,728,923 bp, truncating 94.8% of the fourth exon, and 56.4% of the first of two 3’ UTRs (Table S10). This deletion is present in 194 lines that have a mean protein content 7.9% greater than those with the reference sequence (Table S6), consistent with the high protein haplotype. There is significant overlap between the trinucleotide insertions and the 304 bp deletion: 152 of the 194 individuals with the 304 bp deletion possess a homozygous trinucleotide insertion, and all but one individual with a homozygous trinucleotide insertion had the 304 bp deletion (Table S6). The 304 bp deletion is linked to the GWAS-SNP with R^2^ = 0.918 and D’ = 0.958, and it is most tightly linked with the SNP at position 31,680,574 bp (R^2^ = 0.952, D’ = 0.984). The start position for the 304 bp deletion, 31,728,619 in Wm82.a4.v1 (Valliyodan et al. 2019), coincides with a smaller insertion in some lines (Table S6). The most common insertion at this site is 25 bp, which appears exclusively in 30 of the 194 individuals with the 304 bp deletion (μProtein = 48.29; Table S6).

No significant (MAF > 0.05) InDel sites were identified in the syntenic cqProt-003 region from a Korean collection of 855 soybean individuals (Kim et al. 2021). However, the InDel data available from a recent soybean pangenome (Liu et al. 2020) suggests additional variation may be present in the *20G085100* genic region (annotated as ZH13_20G075300; Table S11). This dataset similarly contains trinucleotides with 3-8 ‘CAA’ repeats across 5.3% of individuals, however they do not identify the remaining six other rare alternate alleles (Table 1, S11). Beyond the trinucleotide tandem repeats, Liu et al.’s data presents similar structural variation in the genic region of *20G085100* (Table S11). The W05, W01, W02, W03 and SoyC12 accessions contain a 318 bp (302 bp for W03) deletion relative to ZH13 and WM82 cultivars, significantly truncating the upstream portion of the fourth exon for *ZH13_20G075300/Wm82_20G085100* (Table S10-11). Individuals with 318 bp deletions in the Liu et al dataset are present in our data with 304 bp deletions; however, no deletion in the *20G085100* gene was identified for W05 using the same genomic data from Liu et al aligned to the Wm82.v1.a4 reference.

## Discussion

Whole genome resequencing datasets from global domesticated and wild soybean populations have provided an opportunity for the detailed characterisation of the cqProt-003 protein associated region, to identify patterns of variation and identify candidate causative variants. We have refined the genomic interval underlying significant protein and oil variability to a 173 kb region. The region defined in our study sits in the centre of a broader 550 kb ‘fourth block’ previously identified using USDA 50k-SNP chip genotyping (Bandillo et al. 2015), however the peak SNP in that block is approximately 400 kb downstream from the GWAS-SNP identified in our study. A GWAS conducted on 279 Chinese soybean cultivars defined a 960 Kb block on chromosome 20 associated with oil (Cao et al. 2017). This seed oil QTL region starts 7 kb upstream of the end of the 173 kb region associated with protein we report (Cao et al. 2017). More recently, an 839 kb linkage block identified in a GWAS of seed protein, oil and amino acids using USDA accessions from maturity groups I to IV is more distant, located 1.5 Mb downstream from our region (Lee et al. 2019). The strong relationship between oil and protein in this region is apparent, indicating that the primary utility of the cqProt-003 locus is for breeders who are willing to forego oil content for higher seed protein, reflecting increasing demand for plant protein for human consumption. The inverse effect could be explained in part by the competing metabolic demands of protein and oil synthesis (Popovic et al. 2012), however further dissection of how the cqProt-003 locus directly impacts absolute levels of yield components is required.

Our results benefit from higher SNP density, InDel data, and genome sequence data sufficient for the accurate detection of a broad range of structural variants. This has allowed us to move beyond marker/trait association to detailing shared haplotypes with specific variants linked to trait variation at the cqProt-003 locus. The characterisation of haplotype structures at this locus provides a detailed view of the local landscape of linkage and genomic variation. Our results show that genomic diversity has been significantly reduced by selection for the high oil/low-protein haplotype, and largely lost in modern breeding pools. Genomic variants are in high linkage throughout the region, which limits the production of novel combinations, and most of the linked clusters of variants (M02-M06) appear to display variation only when the alternate alleles are present in the candidate marker group that is most strongly associated with protein content (M01). The lack of variation in individuals containing the Williams82-like alleles for the M01 group appears to be the result of a strong domestication bottleneck for the high oil/low-protein phenotype in domesticated lines. Further functional characterization of the variation across the different haplotype populations (B-G) is needed to explore how different allelic combinations modulate the effect of cqProt-003 on seed compositional traits.

We identified two primary variants located in the candidate gene for the high protein phenotype, *20G085100*, which remains largely uncharacterized. We located a 304 bp SV in the fourth exon that is associated with the low-protein phenotype. This represents an insertion in modern lines relative to wild progenitor populations, and this insertion is likely to disrupt the expression and/or function of the *20G085100* gene. Previous research that first identified this deletion (reported 321 bp in size) focused in the structural variation between two lines, PI468916 (HNY.50) and Williams 82, before screening a population of 302 individuals using PCR with markers targeting a CCT domain (Fliege 2019). They conclude that this structural variant is the result of the insertion of a transposon fragment in low protein domesticated populations, rather than a deletion in high protein lines. However, little was known about its genomic context and its distribution across different soybean populations. Discrepancies regarding the size of the SV are likely due to different assemblies, Wm82 compared to ZH13 (Liu et al. 2020), and because the PCR fragment amplified by Fliege et al. included 634 bp of the genomic context which likely included other variation. We uncovered additional variation in coding regions, including a 25 bp insertion at the start of the 304 bp SV that suggests multiple distinct orthologues of *20G085100* could have unique influences on seed composition,

We identified a trinucleotide repeat expansion up to 45 bp in the third exon of *20G085100* that is tightly linked with the high protein phenotype. The low-protein allele includes an absence of all tandem repeats, seen in nearly all domesticated lines though rare in wild progenitor populations. This would have resulted in allelic fixation at the site, removing the capability of repeat expansion. Trinucleotide repeat expansions are highly mutagenic structures that have been associated with degenerative diseases in humans (Nageshwaran and Festenstein 2015), especially when present in coding regions, where they can have significant impacts on protein structure (Figura et al. 2015), such as in the case of Huntington’s disease (Shacham et al. 2019). Trinucleotide repeat expansions have also been implicated in temperature-sensitivity adaptation in Arabidopsis thaliana (Tabib et al. 2016), though remain largely unstudied in the plant kingdom (Zhu et al. 2021). Our finding may represent the first case of causal short tandem repeat variation in a coding region underlying phenotypic effect in soybean. We hypothesize that prior to domestication there may have been balancing selection to maintain a moderate number of trinucleotide repeats that may be needed for a function that results in a high level of protein or oil. The lack of this repeat in many domesticated populations may be a significant loss for the adaptive potential of this gene that likely underpins a large degree of trait variability.

## Conclusion

We have refined the linked region associated with the high protein phenotype, defined haplotype structures within this region, thoroughly examined the high-protein haplotype, assessed the untapped variability within the population with this haplotype and identified likely causal genomic candidates. The variants in the high-protein haplotype act as high confidence markers for the high protein haplotype, which can support low-cost genomic inference of the cqProt-003 trait. The causal candidates, trinucleotide insertions and structural variation in the *20G085100* gene, demand validation for impacts on plant phenotype. Furthermore, a key gap in our understanding still remains regarding the role in seed morphology and development of the additional proteins produced by individuals with the high-protein haplotype. A deeper understanding of the functional pathways involving the *20G085100* gene could open the door to further optimization of the cqProt-003 locus for agronomic gain using gene editing technology such as CRISPR. These findings draw attention to the lack of diversity in modern breeding lines at this important locus for seed composition, and the potential to exploit the natural variability remaining in exotic landraces and wild populations (haplotype populations B-G) to provide breeders with additional tools for producing protein rich soy in a world with increasing nutritional demands.

## Data availability

All code generated for this study is available at https://github.com/JacobIMarsh/cqProt-003.

All data generated for this study is available at the UWA research repository (https://www.appliedbioinformatics.com.au/index.php/Soybean) including VCF files for raw, filtered, imputed SNP and filtered InDels recoded as biallelic sites, as well as files used as input for HaplotypeMiner.

JBrowse (Buels et al. 2016) genome visualization of the 40 variants identified in the cqProt-003 high-protein haplotype can be accessed at http://appliedbioinformatics.com.au/soybean_protein/.

## Acknowledgements

The authors thank Australian Research Council for funding support through projects DP200100762, DP210100296, LP160100030 and DE210100398. This work was supported by resources provided by the Pawsey Supercomputing Centre with funding from the Australian Government and the Government of Western Australia.

## Author information

### Contributions

DE supervised the overall project; JM conceived the research and wrote the manuscript with input from HH, PEB, BV and HTN; JIM and HH conducted the bioinformatics analysis, with contributions from JP; JP, PEB, JB and DE contributed to the editing of the manuscript.

### Ethics declarations

#### Conflict of interest

The authors declare no conflict of interests.

## Notes

https://www.appliedbioinformatics.com.au/index.php?title=Soybean

### Competing Interest Statement

The authors have declared no competing interest.

