## Supplemental Figures for "Haplotype mapping uncovers unexplored variation in wild and domesticated soybean at the major protein locus cqProt-003"

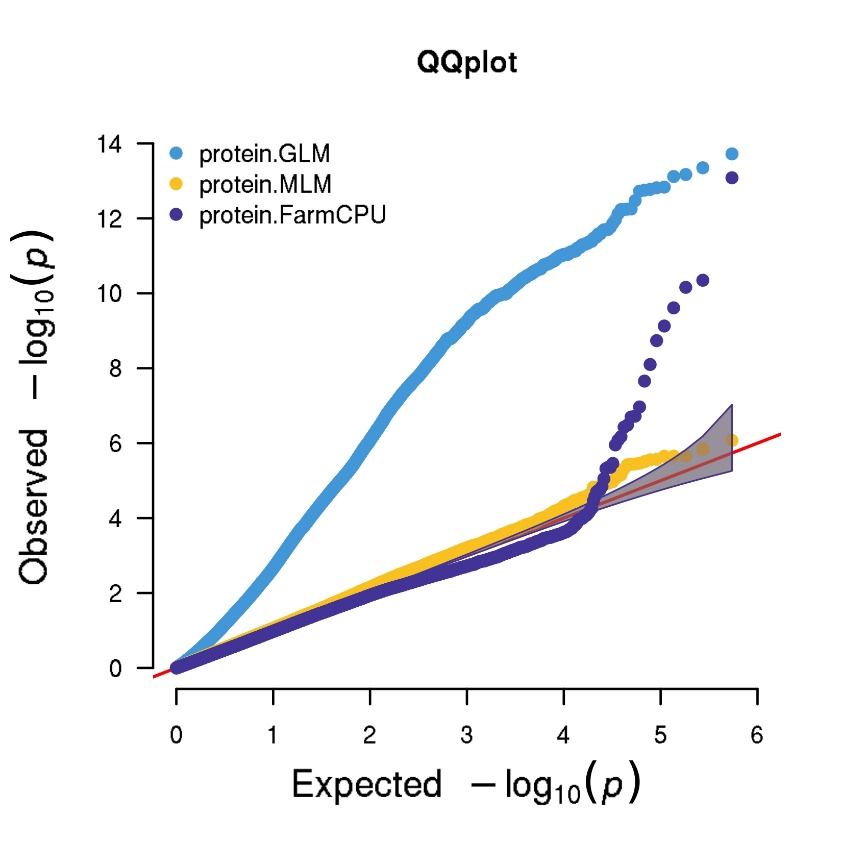

**Figure S1.** **Multitrait QQ plots of GLM, MLM and FarmCPU for protein GWAS using unimputed SNPs** filtered only for quality.

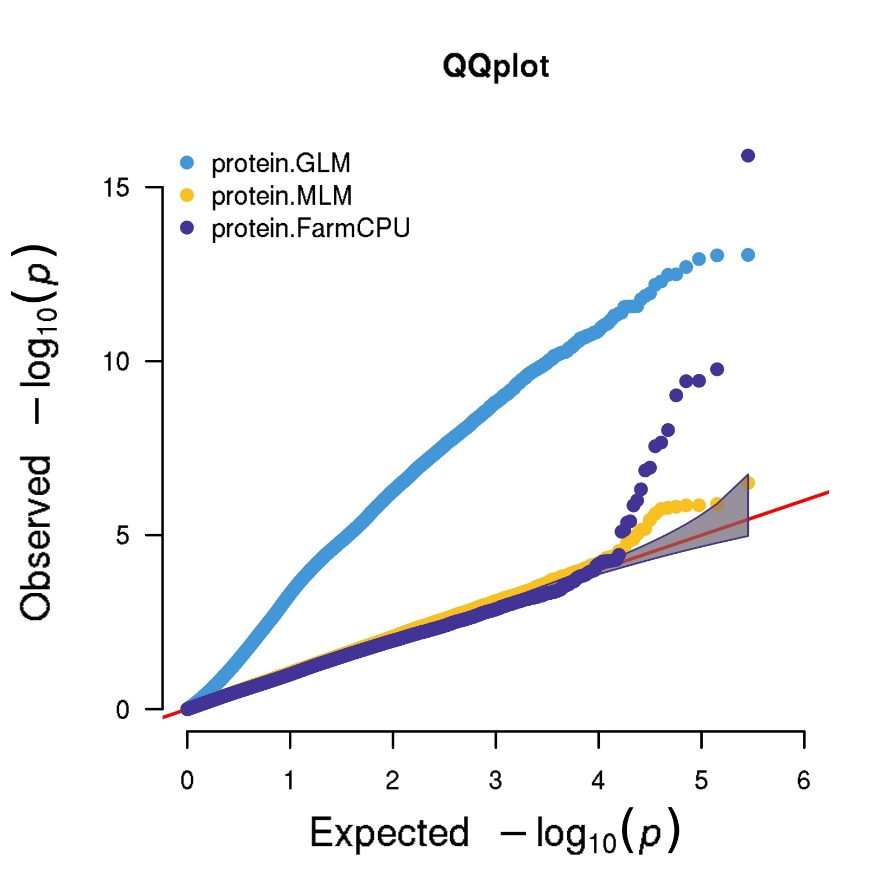

**Figure S2.** **Multitrait QQ plots of GLM, MLM and FarmCPU for protein GWAS using unimputed SNPs and reformatted InDels** filtered for minor allele frequency and missingness.

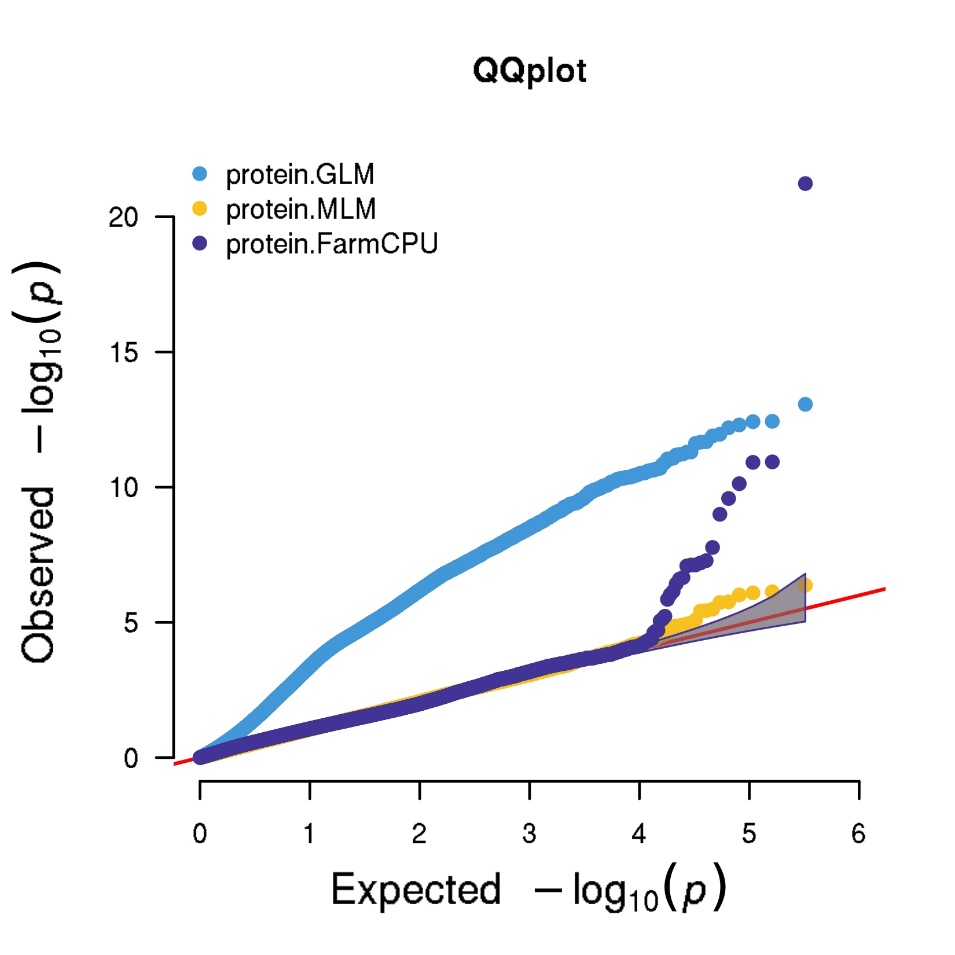

**Figure S3.** **Multitrait QQ plots of GLM, MLM and FarmCPU for protein GWAS using imputed SNPs** filtered for minor allele frequency and missingness.

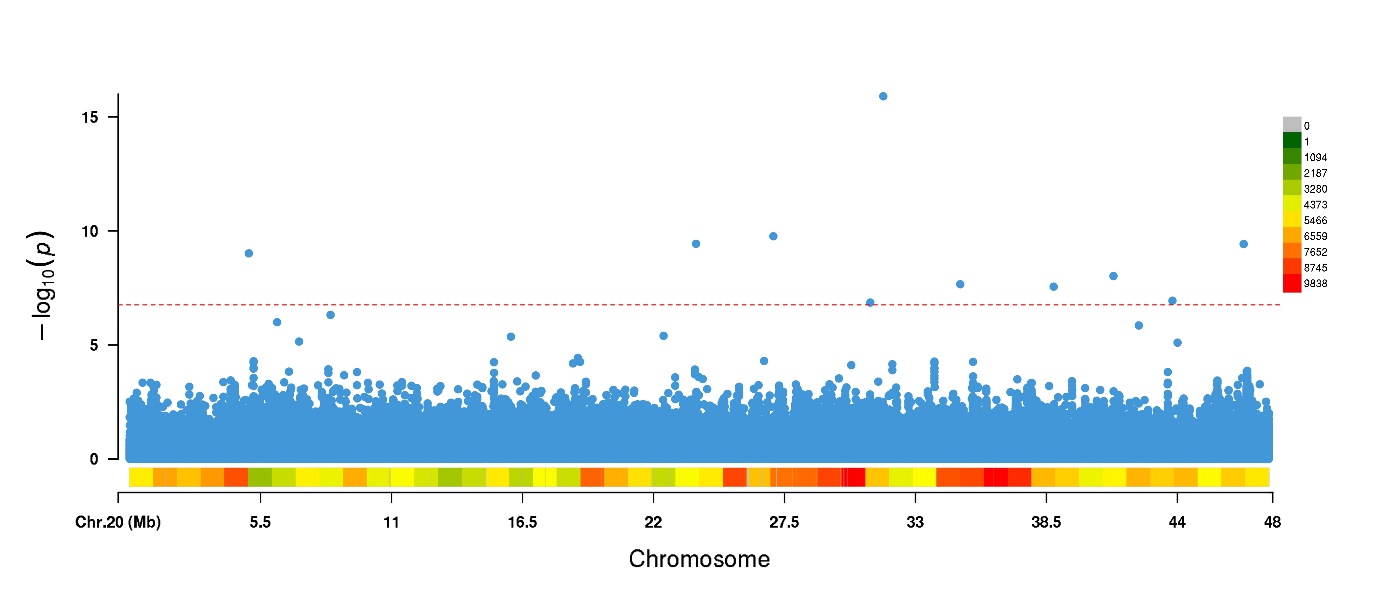

**Figure S4.** **Manhattan plot for protein FarmCPU GWAS** on chromosome 20 using unimputed SNPs filtered for minor allele frequency and missingness.

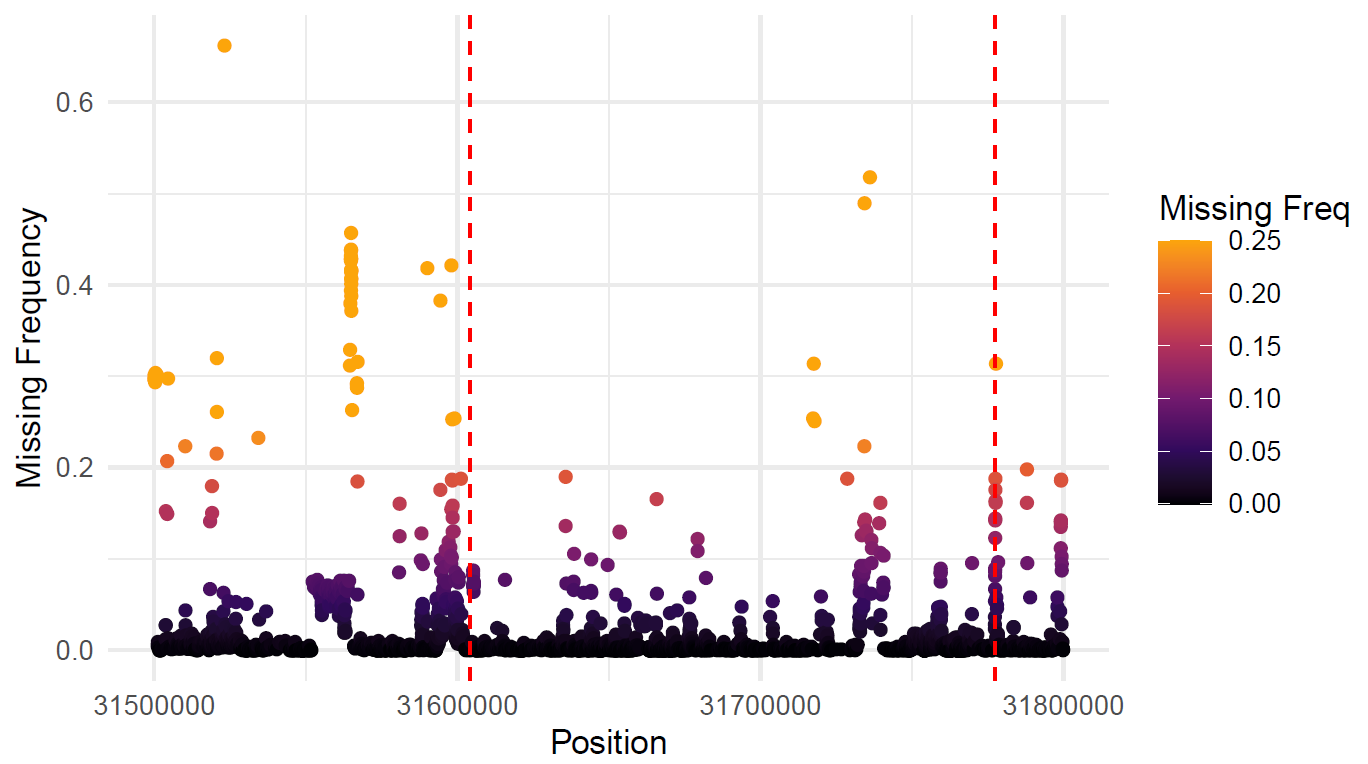

**Figure S5. Frequency of missingness by site around cqProt-003 region.** Variants filtered for minor allele frequency > 0.01. Dashed red lines delimit the 173kb region defined in this study. Each dot represents a SNP locus.

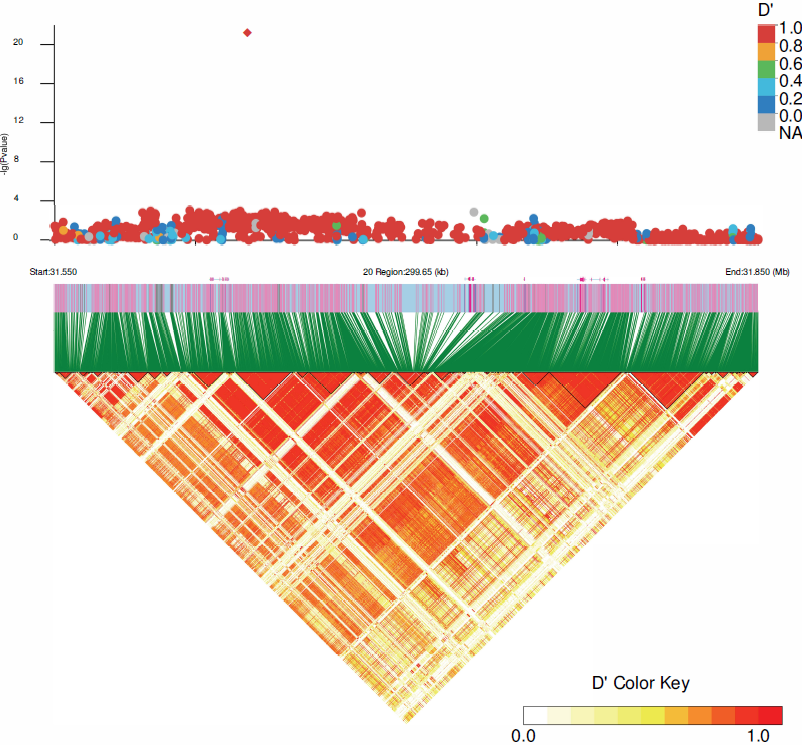

**Figure S6. Linkage disequilibrium of D' between SNPs across a 300Kb region surrounding the significant GWAS-SNP** (red diamond). The thin black lines in the heatmap delineate linkage blocks defined through the confidence interval approach (Purcell et al. 2007).

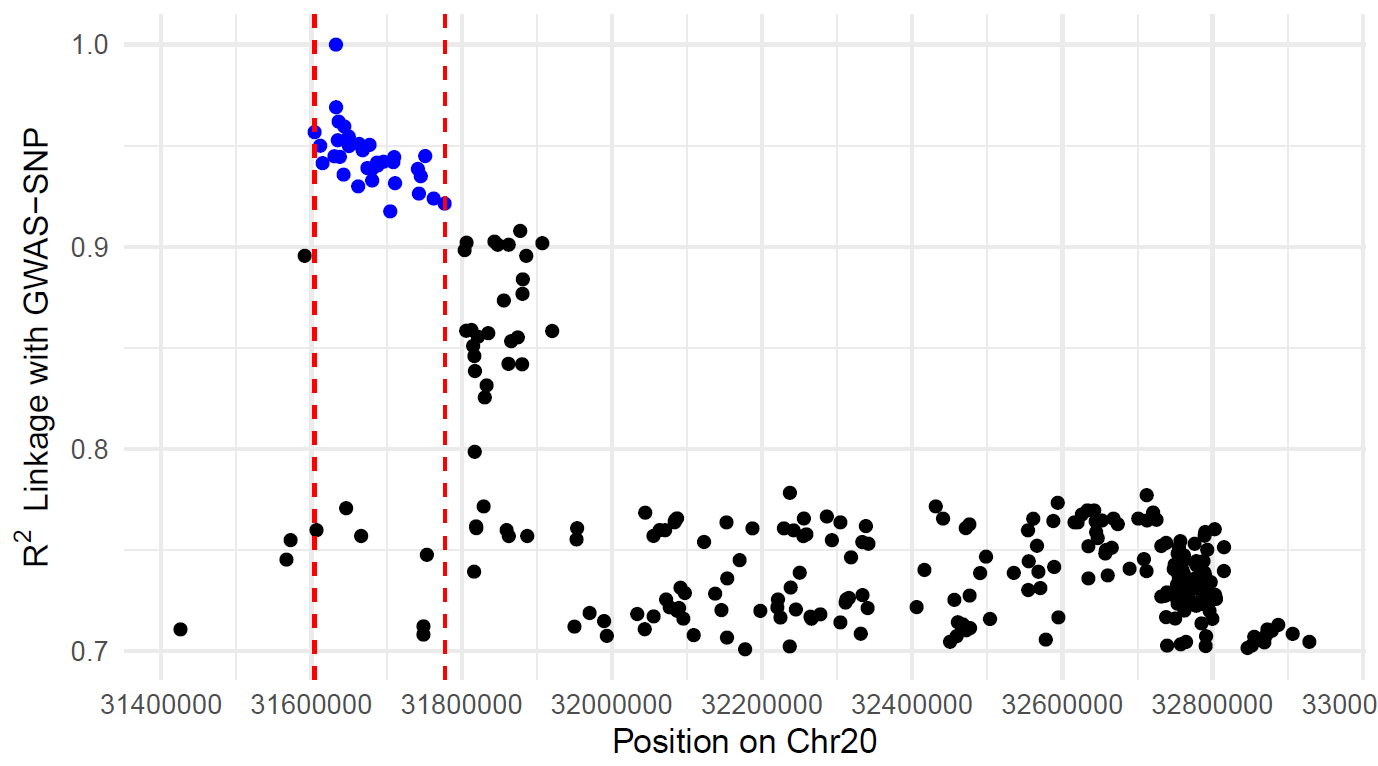

**Figure S7. Linkage (R^2^) of small variants with GWAS-SNP,** by position. Blue points indicate SNP variants included in the M02 marker group (Table S5) and final high protein haplotype (Table S9), dashed red lines delimit the 173kb region.

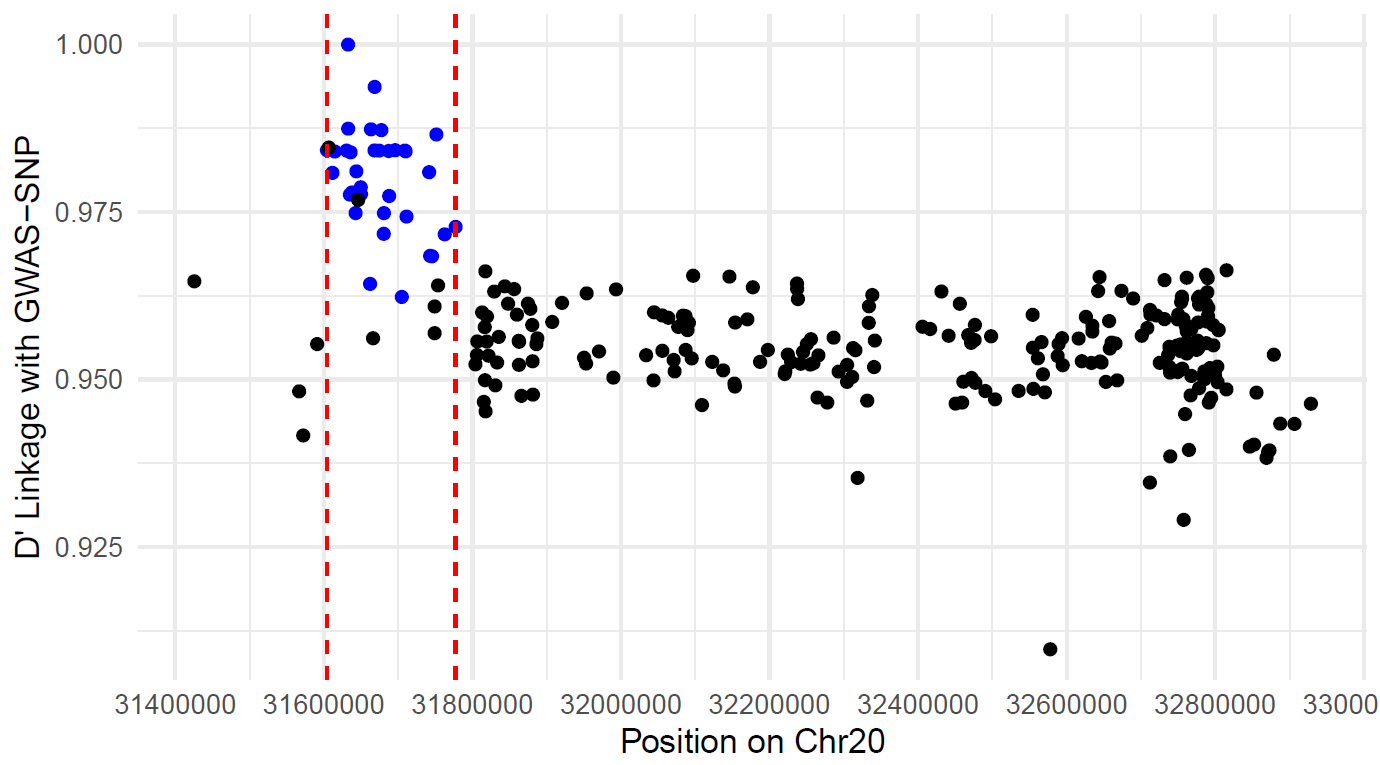

**Figure S8. Linkage (D')** **of small variants with GWAS-SNP,** by position. Blue points indicate SNP variants included in the M02 marker group (Table S5) and final high protein haplotype (Table S9), dashed red lines delimit the 173kb region.

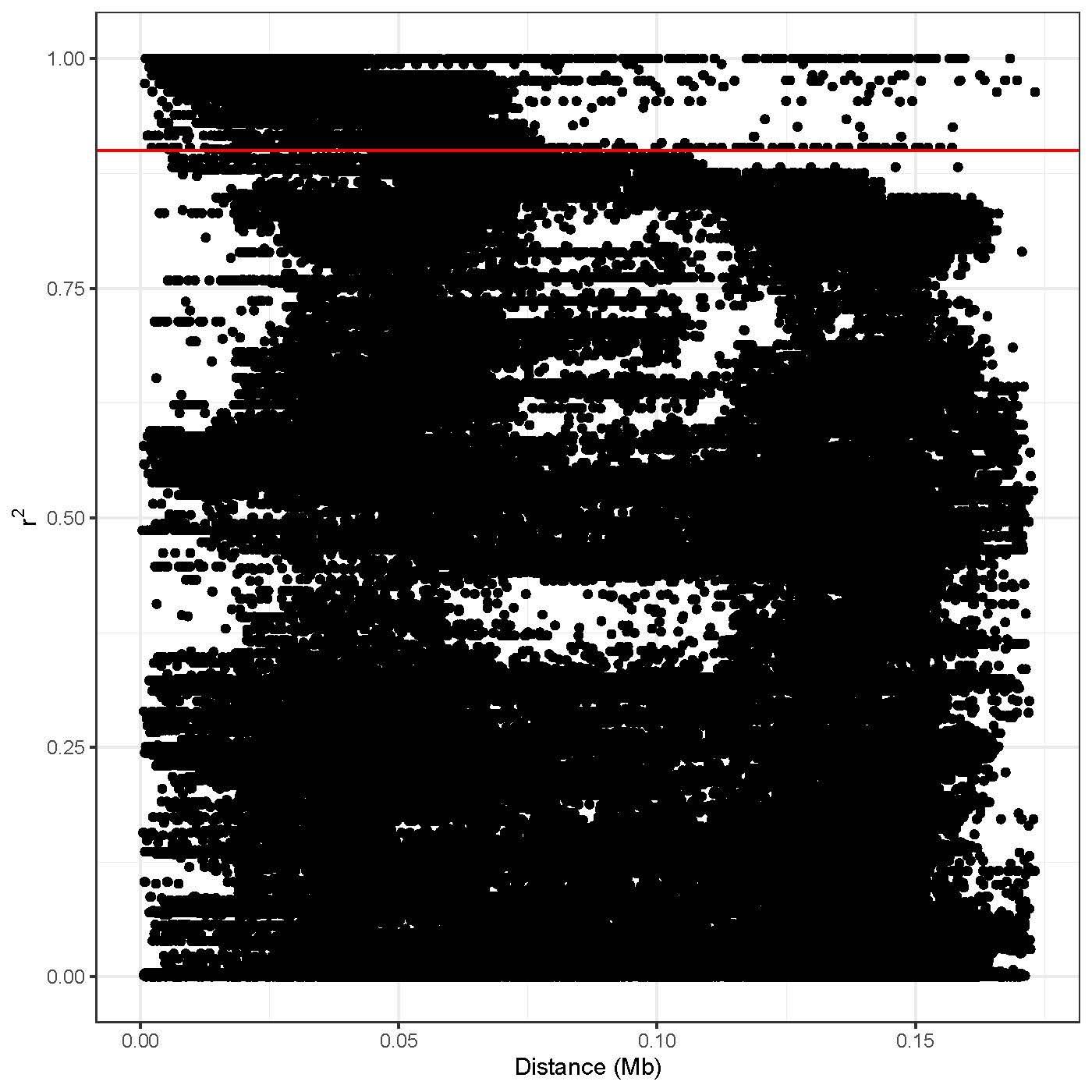

**Figure S9. Estimation of linkage disequilibrium (R^2^) between pairs of markers across the 173kb region as a function of pairwise distance** taken directly from HaplotypeMiner (Tardivel et al. 2018) output**.** The horizontal red line corresponds to marker_indepenence_threshold R^2^>0.9.

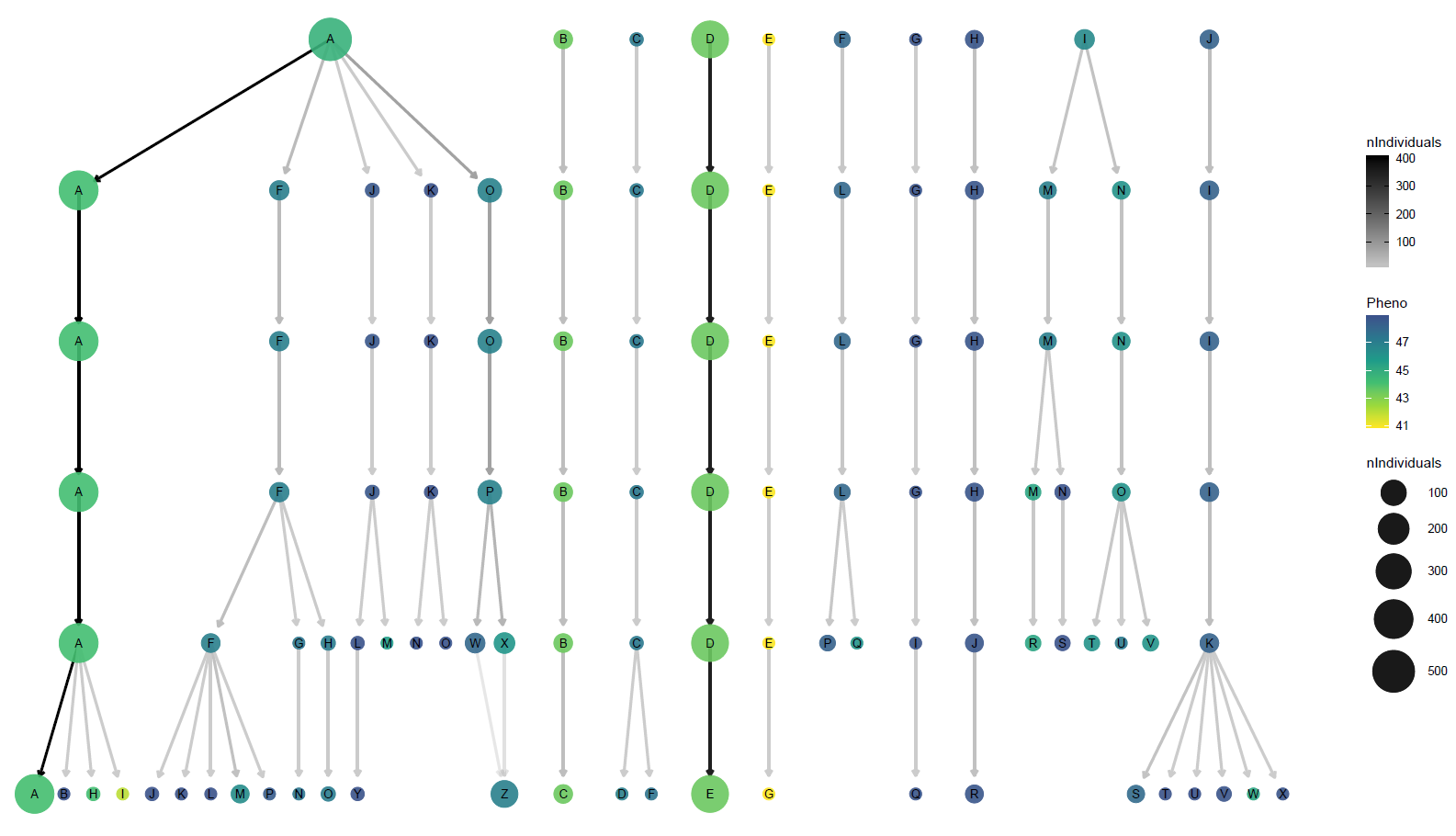

**Figure S10: Stability of clusters of individuals who share haplotype combinations at different cluster thresholds (CT)**. CT= 0.5 at the top, each lower row represents a decrease in CT of 0.1, with CT = 1.0 at the bottom; the chosen CT of 0.6 is highlighted by the red circle. N.B. this includes clusters removed due to redundancy, thus the letters for haplotype populations do directly correspond to those reported in the manuscript.

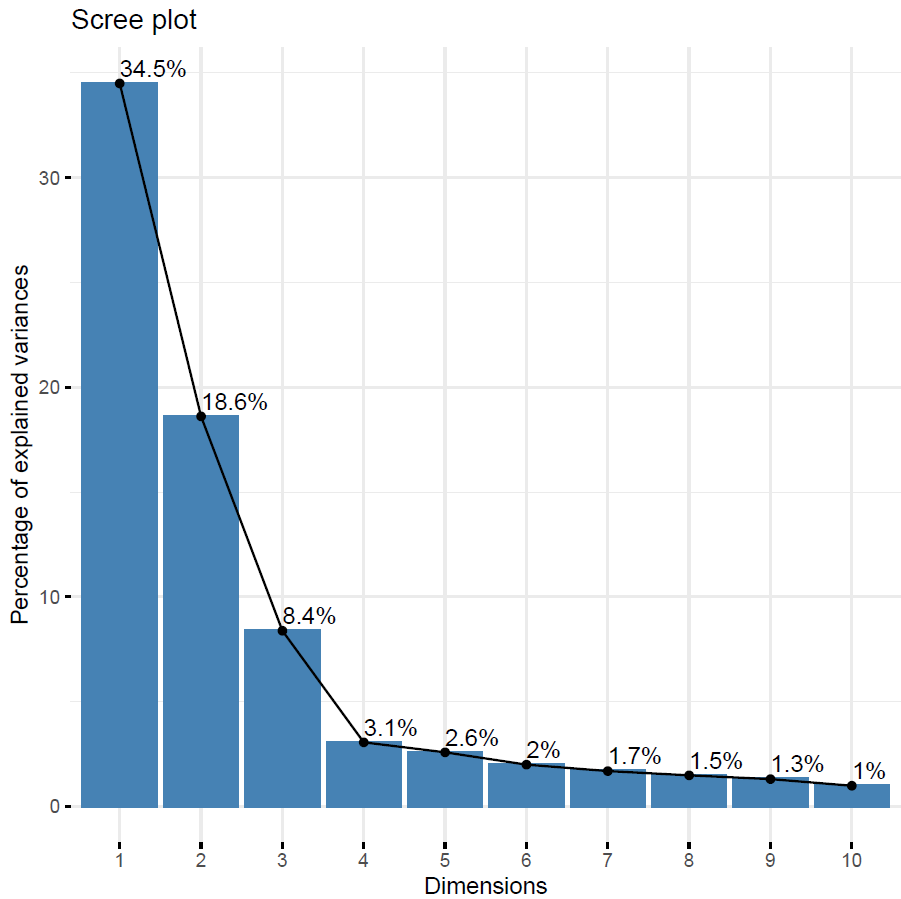

**Figure S11: Scree plot displaying proportion of total genetic variance between individuals explained by principal components.** PCA was run across individuals using SNPs from within the 173kb region only.

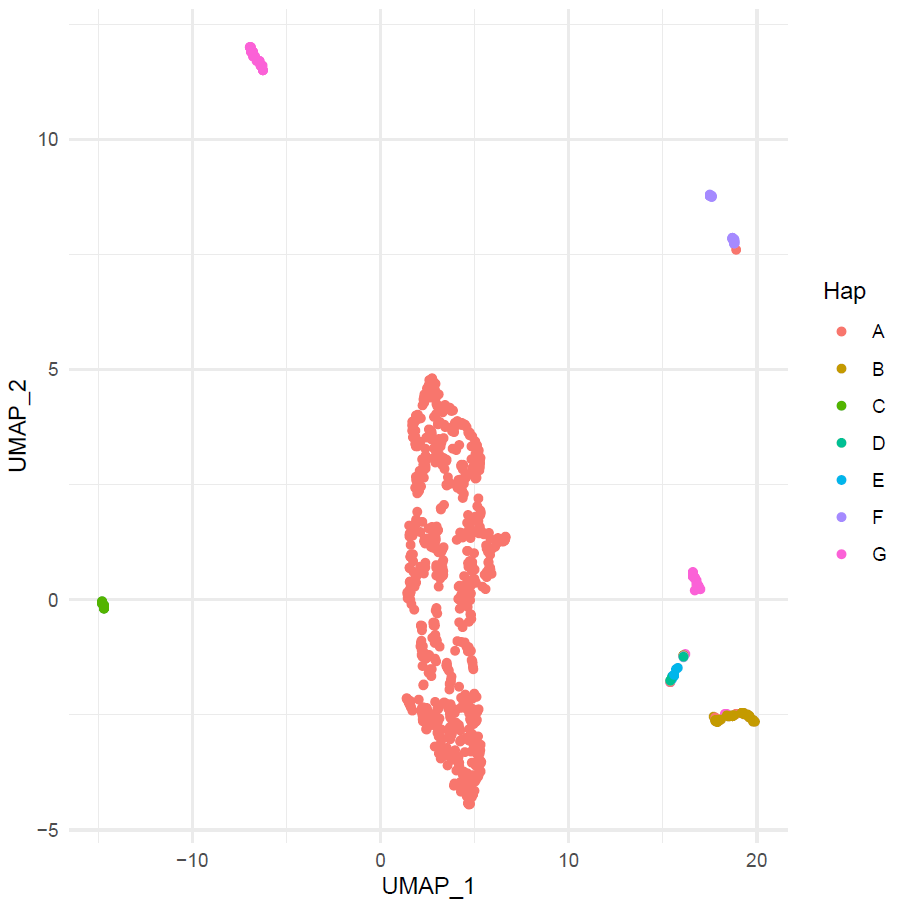

**Figure S12: Dimensionally reduced projection (UMAP) for assessing segmentation between haplotype groups (A-G).** Each dot represents an individual, clustered based on genetic similarity in the 173kb region. The Hap A individuals that are not located in the central cluster can be explained as being 6 atypical A haplotype individuals (Table S6): HN039 is wild and heterozygous at GWAS-SNP; HN092 and SRR1533182 are heterozygous for trinucleotide insertions at 31,727,019 bp; PI507638 is wild and has the 304bp deletion at 31,728,619 bp; PI378696B and PI378696A are wild and contain homozygous trinucleotide insertions at 31,727,019 bp.

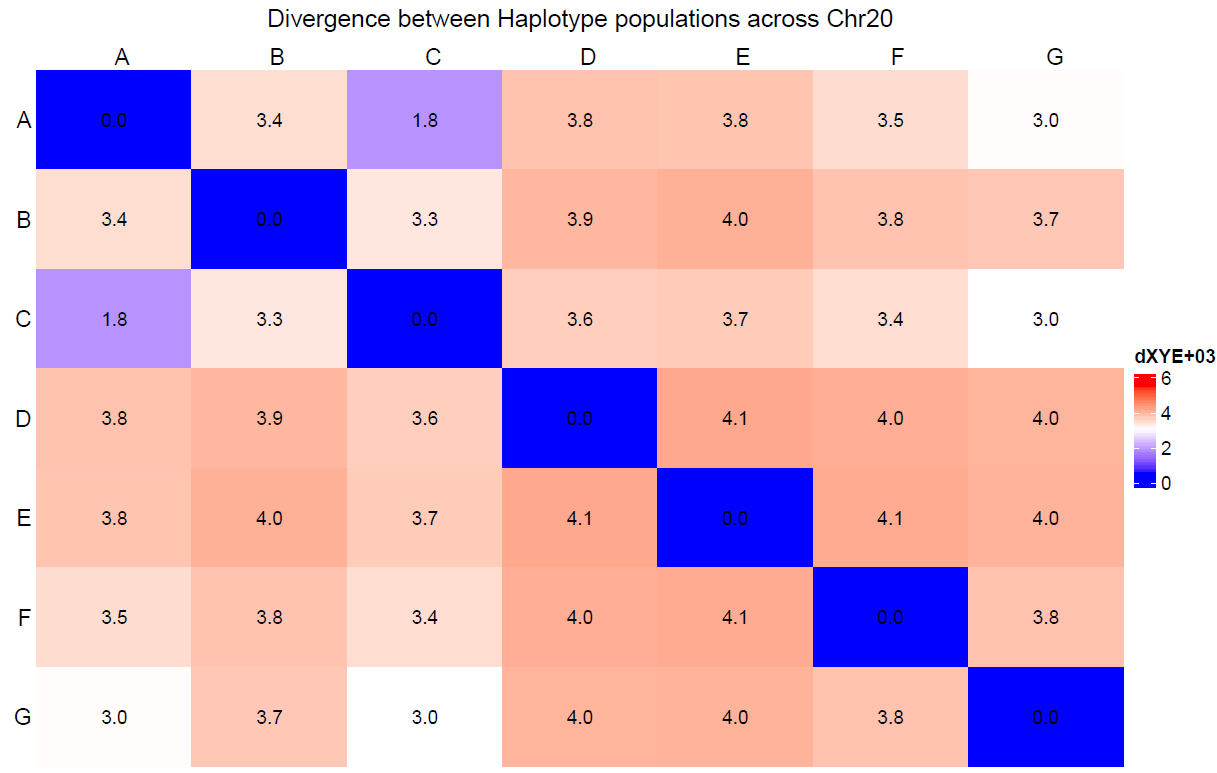

**Figure S13. Divergence (dXY) between haplotype group populations across chromosome 20**. Using filtered, unimputed SNPs.

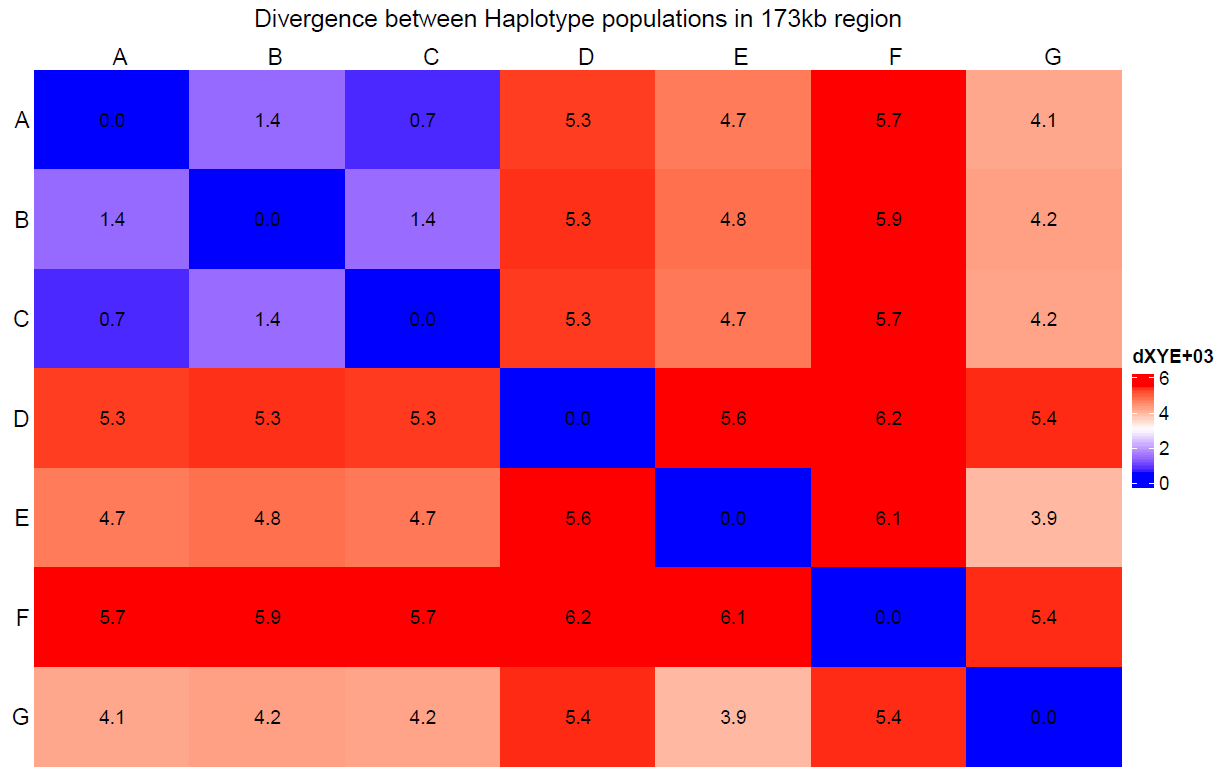

**Figure S14. Divergence (dXY) between haplotype group populations within the 173kb region**. Using filtered, unimputed SNPs.
